# Modeling Symbiont Dynamics and Coral Regulation under Changing Temperatures*

**DOI:** 10.1101/2024.04.10.588659

**Authors:** Jerome Cavailles, Christoph Kuzmics, Martin Grube

**Author notes:** We are grateful to Andrew Baker, Michael Greinecker and the members of the field of excellence initiative COLIBRI at the University of Graz for helpful comments and suggestions.

## Abstract

Corals play an essential role in marine ecosystems by creating protective coastal structures and habitats for marine biodiversity. Their symbiotic relationship with various algal species, where corals supply nitrogen in exchange for carbon products, is vital for their survival. However, with some algal species being temperature sensitive, this vital symbiosis is increasingly threatened by global warming, causing significant symbiont losses, potentially leading to coral bleaching and fatal consequences. Here, we model the optimal regulation of algal populations by corals through nitrogen allocation. Two algal species compete for nitrogen: one is effective in carbon supply and rapid growth, and the other is resilient to temperature increases. Our testable analytical solution identifies the optimal total algal population as a function of the current temperature and symbiont composition. The model also determines the relative abundances of the two algal species based on current and historical temperatures. Our findings are consistent with numerous previous observations and experimental studies. The model clarifies how inter-species competition under varying temperature patterns shapes the composition and dynamics of algal species in coral symbiosis. It also clarifies that bleaching occurs when the relatively efficient algae fail to exchange enough carbon products at high temperatures.

## 1 Introduction

Coral reefs are indispensable to marine ecosystems and human communities alike, offering a myriad of crucial services. They serve as natural barriers that protect coastlines from storms and erosion. Additionally, coral reefs are vibrant habitats for a diverse array of marine species, fostering rich biodiversity. This biodiversity includes fish populations that are vital for the sustenance of millions of people, particularly in vulnerable tropical regions. Beyond their ecological importance, coral reefs contribute billions of dollars to the global economy through fisheries, tourism, and coastal protection [Cesar et al., 2003]. The economic valuation of coral reefs underscores their critical role in environmental health and human prosperity [Woodhead et al., 2019, Eddy et al., 2021]. Unfortunately, the past few decades have seen a dramatic decline in coral cover, driven by anthropogenic climate change [Bellwood et al., 2019]. The Intergovernmental Panel on Climate Change (IPCC) has highlighted the heightened risk to corals, warning of severe implications if current trends continue [Pörtner et al., 2019]. This escalating crisis calls for urgent attention and action to safeguard these ecosystems. At the heart of coral vulnerability is their dependence on a symbiotic relationship with algae, a partnership that is essential for their survival.

Corals maintain a vital symbiotic relationship with algae to meet their energy requirements. These algae perform photosynthesis, producing carbon-rich compounds that serve as the primary energy source for corals [Muscatine, 1990]. In return, corals provide nitrogen and other essential nutrients to the algae, creating a mutualistic exchange vital for both organisms [Yellowlees et al., 2008]. This symbiosis significantly influences coral fitness and health, as the energy derived from algae is fundamental to coral growth and reproduction. The regulation of algal density is a key adaptive mechanism that corals employ to maintain optimal symbiotic functionality under varying environmental conditions. Under harsh environmental conditions, this equilibrium can be upset, causing disruptions in the symbiotic relationship.

A massive expulsion of algal symbionts by corals is known as *bleaching*. This phenomenon causes corals to turn white as they lose the algae that gives them color. Since the late 20th century, the frequency [Hughes et al., 2018] and severity of bleaching events have increased significantly, particularly during periods of elevated temperatures [Gates et al., 1992]. This alarming trend has raised concerns about the long-term health and survival of coral reefs globally. Buddemeier and Fautin [1993] proposed that bleaching facilitates the recolonization of the host with a different partner. This hypothesis suggests that, following a bleaching event, corals could be recolonized by different algal species, potentially those better suited to the new environmental conditions. Supporting this idea, several studies have documented changes in symbiont composition during and after bleaching events Silverstein et al. [2015], Baker [2003], Quigley et al. [2022]. Furthermore, Baker [2001] described bleaching as a “risky opportunity” for corals, presenting a chance to establish symbiotic relationships with more temperature-tolerant algae. Observations by Guest et al. [2012] showing that corals with a history of thermal stress are more resilient to subsequent bleach-ing events, are consistent with this theory. For a comprehensive explanation of the adaptive bleaching hypothesis and an extensive review of its fundamental features, including various tested assumptions and specific mechanisms, refer to Buddemeier et al. [2004]. These adaptive mechanisms highlight the dynamic nature of coral-algal symbiosis, revealing how corals might mitigate the impacts of environmental stressors through flexible and evolving partnerships.

Corals can host various lineages of algal species throughout their lives, as revealed by DNA sequence analyses [Mieog et al., 2007]. For detailed taxonomic classifications, refer to LaJeunesse et al. [2018]. Some studies have shown that corals may simultaneously harbor multiple algal species [Baker, 2003, Little et al., 2004, Quigley et al., 2014]. Typically, one symbiont dominates while the others remain in the background [Little et al., 2004]. Flexibility in symbiotic relationships can enhance the overall resilience of coral-algal symbiosis. Different algal types have varying physiological demands and can adapt to different ecological situations, which can be advantageous in fluctuating environments [Grégoire et al., 2017]. Moreover, Herrera et al. [2021] found that temperature can override partner specificity in the establishment of cnidarian symbiosis, suggesting that environmental factors can significantly influence symbiotic dynamics.

To understand the intricate dynamics of coral-algae symbiosis, and their responses to environmental stressors, developments in mathematical modeling are essential. Several models have been proposed to predict bleaching frequency in the coming decades [Weijerman et al., 2015], yet there remains a need for improved models that link environmental history with bleaching patterns [Van Woesik et al., 2022]. Ware et al. [1996] modeled the adaptive bleaching hypothesis with competing algae but did not include the active control by the host in regulating algal populations through nutrient allocation. Muller et al. [2009] developed a dynamic bioenergetic model (DEB) of the coral-algal symbiosis. This approach was utilized by Cunning et al. [2017] to describe bleaching as an alternative stable state and more recently by Kaare-Rasmussen et al. [2023] to fit the model with data from the anemone *Exaiptasia pallida*. Brown et al. [2022] developed a model that explicitly considers the shuffling between sensitive and tolerant symbionts and the occurrence of bleaching. Using a specific setting, van Woesik et al. [2010] modeled how corals shuffle between different algal species. Baskett et al. [2009], and later Logan et al. [2021], compared the importance of symbiont shuffling with symbiont evolution. Various models have also explored the evolution of the interaction between corals and algae. For instance, Day et al. [2008] modeled the evolution of bleaching resistance, while Raharinirina et al. [2017] investigated the evolution of symbiotic investment in corals. In a subsequent study, Raharinirina et al. [2022] used a similar model to predict the consequences of different climate scenarios. To incorporate corals into a broader framework, more complex multi-scale models have been developed. Baskett et al. [2010] proposed a five-layer model encompassing locations, species, coral size structure, symbiont populations, and symbiont genetics. Generally, models have been developed to formalize the competition between corals and other organisms, such as macro-algae, to predict the impact of climate change on coral ecosystems [Fabina et al., 2015, Mumby et al., 2007, Baskett et al., 2014].

As an alternative to previous approaches, we have developed a simple and analytically solvable model that incorporates the active role of coral in regulating symbiosis. This model aims to provide a deeper understanding of the key mechanisms driving changes in symbiotic composition and the resulting susceptibility to bleaching. Additionally, the predictions of our model can be statistically tested, offering a practical tool for researchers. As suggested by Weijerman et al. [2015] minimal models can be useful to predict the shape of disturbance response. Our model uses only a few parameters yet effectively captures various observed features, including dynamic changes in algae within coral symbiosis and the severity of bleaching relative to historical temperatures. We begin by introducing a stylized version of the model, demonstrating its ability to explain several empirical observations. We then generalize the model to account for a broader range of empirical data and propose various testable hypotheses. The core principle of our model is the coral’s ability to establish symbiosis with different algal species, allowing rapid adaptation to diverse environmental conditions, particularly higher sea temperatures. This adaptability is evolutionarily advantageous as long as environmental changes do not occur too rapidly. Our model shows that competition between algal species and their respective growth rates in different environments can explain how prolonged warm temperatures favor temperature-tolerant algae. In the same environment, one algal species will always out-compete the others, while the less competitive algae remain in the background.

## 2 The basic model

In this paper, we consider an individual coral living in an environment with fluctuating temperatures and engaged in a symbiotic relationship with algae. The coral exerts active control over the symbiosis by regulating the amount of nutrients (nitrogen) it supplies to the algae. This regulatory mechanism is crucial as it directly impacts the population dynamics of the algae, thereby influencing the overall health and stability of the coral-algae symbiosis. By adjusting nutrient allocation, the coral can adapt to changing environmental conditions, ensuring its survival and optimizing its energy acquisition from the algae.

### The Environment

Time is modeled as discrete and is noted by *t* = 0, 1, 2, ^1^ The environmental state at time *t* is represented by *θ*_*t*_, which can take one of two values: *θ*_*t*_ = 0 indicating an unfavorable (warm) environment, and *θ*_*t*_ = 1 representing a favorable (cool) environment. We extend this binary simplification to an arbitrary number of states in Section 5. All results presented in this manuscript hold for any sequence of *θ*_*t*_, with exceptions detailed in Appendix E. The sequence of *θ*_*t*_ (a series of zeros and ones) can be generated through any method. For instance, *θ*_*t*_ can follow a stochastic process such as a Markov chain, transitioning between the two temperature states.^2^

### Algae Dynamics

Corals live in symbiosis with algae. The coral supplies nutrients to the entire algal population, which in turn provides the coral with carbon products fixed through photosynthesis [Yellowlees et al., 2008]. Corals engage with different algal species simultaneously. Similar to Logan et al. [2021], we limit our model to two key species: Cladocopium and Durusdinium. Cladocopium is more efficient at carbon fixation [Ezzat et al., 2017, Baker et al., 2013b], whereas Durusdinium exhibits greater resilience to elevated temperatures [LaJeunesse et al., 2010, Stat and Gates, 2011, Baker et al., 2013b]. ^3^ At each time step, the density (or abundance) of Cladocopium is denoted by *y*_*t*_, and the density of Durusdinium by *z*_*t*_. These two algal species compete for the nutrients provided by the coral. This competition is reflected in their growth rates, which are influenced by the state of the environment. The relevant quantity for this competition is the relative growth rate of the two algal species (the growth rate of the efficient Cladocopium divided by the growth rate of the resilient Durusdinium). We denote this relative growth rate by *ρ*(*θ*_*t*_) and assume that:

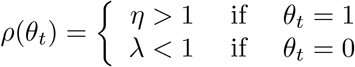

These assumptions are based on empirical evidence indicating that the efficient algae grows faster than the resilient one in a cool environment (*η >* 1) [Little et al., 2004, Wang et al., 2022], but slower in a warm environment (*λ <* 1) [Chen et al., 2005, Fabricius et al., 2004, LaJeunesse et al., 2009, Wang et al., 2022].

We consider a simple model of competition between the two algal species [Krueger et al., 2020, McIlroy et al., 2019, 2020]. Specifically, our model focuses on the competition among symbionts that are already present (even if in minimal quantities) within the coral host. Observations have shown that *shuffling* a change in the proportion of algae species already associated with the host is more significant in coral symbiosis than *switching*, which involves establishing new symbiotic relationships with different algae species [Berkelmans and Van Oppen, 2006]. Nevertheless, we evaluate the influence of symbiont switching in Appendix D. Additionally, it has been demonstrated that shuffling plays a more critical role than symbiont evolution in adapting to environmental changes [Logan et al., 2021]. The dynamics of the two algal species are described by the following equations:

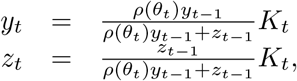

where *K*_*t*_ represents the amount of nutrients available to the algae at time *t*. It can be verified that the total algal population at time *t* equals *K* _*t*_. This quantity serves as the controlled variable for the coral, meaning that the coral can choose the value *K*_*t*_ to maximize its fitness. We assume that the algae exhibit a growth rate that surpasses the coral’s ability to promptly adjust its nutrient provision. Consequently, at each time step, even with changes in nutrient availability (*K*_*t*_), the algae can rapidly reach their carrying capacity, efficiently ut ilizing al l the nutrients provided by the coral by the end of the time step. In our model, the composition of each algal species is determined by their respective growth rates. The ratio of each algal species changes proportionally to the ratio of their growth rates. For instance, if the growth-rate ratio between the efficient an d resilient algae is 2, the relative proportion of efficient al gae wi ll do uble fr om on e point in time to the next. If the growth rates are identical, the species proportions remain constant, though the total population may vary.

### Coral Strategies

Research has demonstrated that corals can actively regulate the amount of nutrients they provide to their algal symbionts [Morris et al., 2019, Falkowski et al., 1993, Rees, 1991, Yellowlees et al., 2008, Cui et al., 2018]. Specifically, c orals c ontrol n itrogen l evels [Thies e t a l., 2 022, Rädecker et al., 2023] in species such as Aiptasia [Cui et al., 2019], Exaiptasia pallida [Xiang et al., 2020], and Stylophora pistillata [Cui et al., 2022, Krueger et al., 2020].^4^ In our model, we assume that the coral can control the amount of nutrients *K*_*t*_ provided to the algal population at each time step. When determining the quantity *K*_*t*_, the coral can perceive the current environmental state *θ*_*t*_ and the densities of the two algal species, *y*_*t*−1_ and *z*_*t*−1_, from the previous period. This timing reflects the notion that algae grow faster than the coral can adapt its nutrient provision.

### Coral Payoffs

The coral optimizes a tradeoff between receiving benefits (carbon) from the algae and providing costly nutrients (nitrogen) to the algae. Formally, at each time step *t*, the coral chooses *K*_*t*_ to maximize its *payoff*

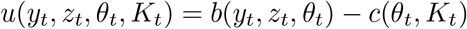

In the basic model, we select the following benefit and cost functions to capture essential empirical features. We consider a generalization in Appendix C. The benefit f unction i s d esigned t o b e c oncave, r eflecting di minishing re turns as the algal population increases, while the cost function is linear, representing a constant marginal cost of nutrient provision. The parameter *α <* 1 represents the efficiency trade-off between the two algae species [Sproles et al., 2020].

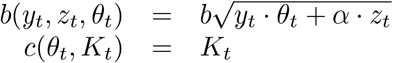

These functional forms are supported by empirical evidence. According to Kemp et al. [2023], under ambient conditions, coral colonies hosting either Durusdinium or Cladocopium exhibited similar nutrient exchange rates. However, during heat exposure, colonies with Durusdinium experienced less physiological stress and maintained higher carbon assimilation and nutrient transfer compared to those with Cladocopium. The concave benefit f unction i s consistent with findings that carbon acquisition and translocation rates p er symbiont cell decrease as symbiont density increases [Pupier et al., 2019, Rossi et al., 2018]. This aligns with the model proposed by Cunning and Baker [2014] and used by Raharinirina et al. [2017], where benefits diminish as algal density grows.

At each time step, the sea temperature is given by *θ*_*t*_. Knowing the current state *θ*_*t*_, the coral selects the optimal *K*_*t*_ based on the previous densities of the two algal species, *y*_*t*−1_ and *z*_*t*−1_. After one period of algal growth, the payoff *u*(*y*_*t*_, *z*_*t*_, *θ*_*t*_, *K*_*t*_) is realized at the end of the time step.

**Table 1:** Overview of key notation.

| Symbol | Description |
| --- | --- |
| $t$ | time |
| $y_t$ | density of the relatively efficient algae species at time $t$ |
| $z_t$ | density of the temperature-resilient algae species at time $t$ |
| $\sigma_t$ | ratio $y_t/z_t$ |
| $K_t$ | amount of nutrients available to the algae at time $t$ |
| $\theta_t$ | temperature at time $t$ |
| $\eta$ | relative growth rate of the efficient alga species to the resilient one in the cold environment |
| $\lambda$ | relative growth rate of the efficient alga species to the resilient one in the warm environment |
| $\alpha$ | efficiency trade-off between the two algae species |
| $\rho(\theta_t)$ | relative growth rate of the efficient species to the resilient one at temperature $\theta_t$ |
| $u(y_t, z_t, \theta_t, K_t)$ | coral payoffs at time $t$ |
| $b(y_t, z_t, \theta_t)$ | benefits to the coral at time $t$ |
| $c(\theta_t, K_t)$ | cost to the coral at time $t$ |

## 3 Coral optimal nutrient provision

The primary hypothesis of this paper is that evolutionary pressures have shaped coral behavior to act as if they maximize their payoff function, considering the constraints imposed by algal dynamics. At each time step *t*, the coral’s objective is to maximize its payoff by optimally selecting the amount of nutrients *K*_*t*_ to provide. The problem is formulated as:

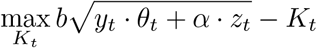

subject to

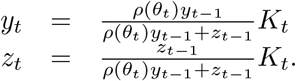

This can be solved by substituting the constraints into the objective function, transforming it into an unconstrained optimization problem,

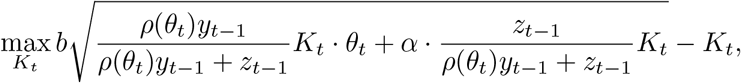

which can be simplified to

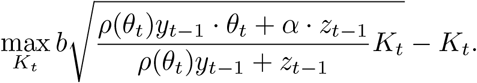

We observe that only the relative density of the two algae species enters this optimization problem. Denoting the relative density by 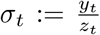 for all *t*, we get

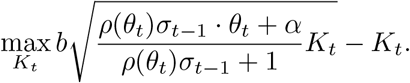

The first-order condition for maximizing the payoff (given the functional form) is obtained by equating the derivative of this objective function to zero.

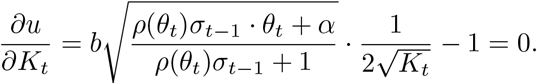

Solving this equation provides the optimal nutrient provision *K*_*t*_ as a function of the environmental state *θ*_*t*_ and the relative abundance of the two algal species from the previous period *σ*_*t*−1_. The solution is given by

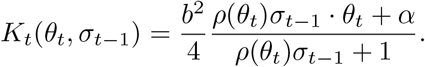

For the two environmental states, we have:

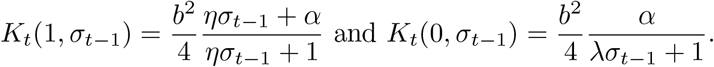

Since the model is analytically solvable, we can analyze its response to changes in parameter values over time. Notably, we can observe how *K*_*t*_, the optimal nutrient provision, reacts to *η*, the relative growth rate of the efficient algae in a cool environment. We find that *K*_*t*_ increases with *η*, but the rate of increase diminishes as *η* becomes larger:

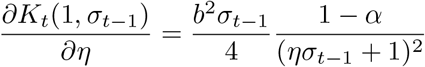

Similarly, we can examine how *K*_*t*_ varies with *α*, representing the relative efficiency of the resilient algae compared to the efficient one. In this case, as *α* increases, *K*_*t*_ also increases, with the change being more pronounced when the resilient algae are more abundant:

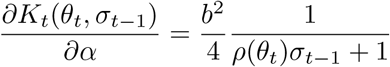

## 4 Model consistency with empirical observations

### Model validation

Figures 2, 3, and 4 compare empirical observations from the literature with the output of our model. The goal is not to parameterize the model for precise quantitative predictions but to qualitatively reproduce observed patterns. Our model, with its minimal parameters, effectively captures the qualitative changes in the abundance of each algal species and the corresponding bleaching severity.

**Figure 1:**
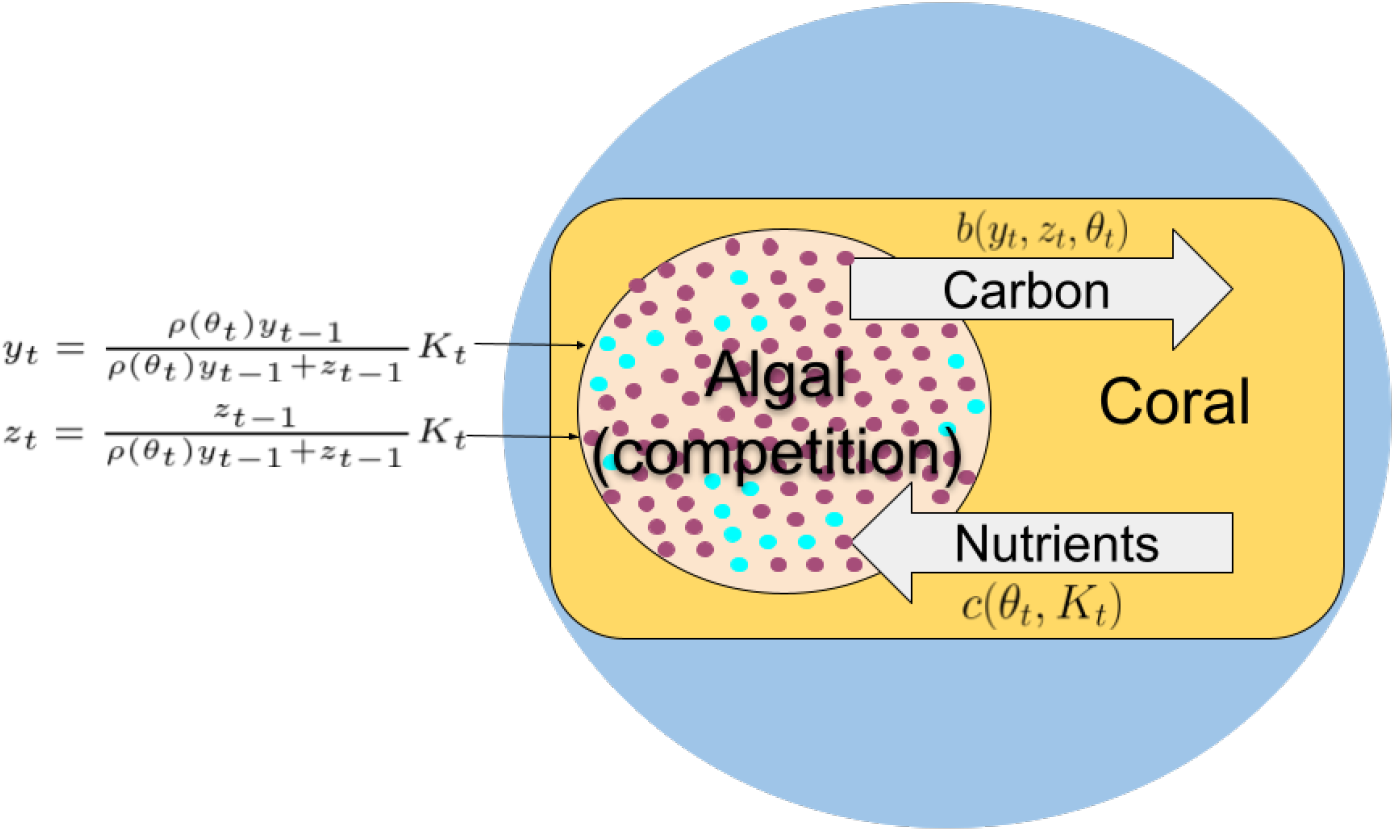
Model Illustration: Corals live in the sea, where the temperature (*θ*_*t*_) fluctuates over time. Within the coral’s cells, two types of algae compete for nutrients. Cladocopium (*y*_*t*_) is relatively efficient in carbon fixation and Durusdinium (*z*_*t*_) is temperature-resilient. The relative growth rate of the efficient algae (*ρ*(*θ*_*t*_)) depends on the current temperature. The coral regulates the nutrient supply, denoted by *K*_*t*_, to control the total algal population. In return, the coral receives carbon products from the population of algae, quantified by *b*(*y*_*t*_, *z*_*t*_, *θ*_*t*_). The coral optimizes its nutrient allocation *K*_*t*_ to maximize its payoff, defined as *u* = *b*(*y*_*t*_, *z*_*t*_, *θ*_*t*_) − *c*(*θ*_*t*_, *K*_*t*_), where the cost of providing nutrients is *c*(*θ*_*t*_, *K*_*t*_) = *K*_*t*_. This regulatory mechanism allows the coral to acclimatize to environmental changes.

**Figure 2:**
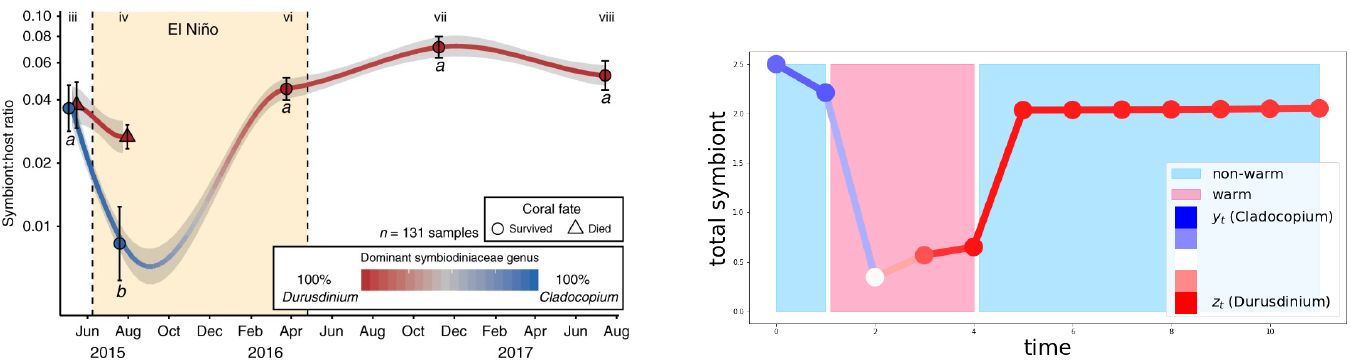
Comparison between observations from Claar et al. [2020] (Figure reproduced unchanged with the kind permission of the authors, CC) and the output of our model. The model reproduces the shuffling of symbionts during the warm period (El Niño), shifting from a predominance of Cladocopium (blue) to Durusdinium (red). Additionally, the model shows a decrease in the total symbiont abundance during the warm period.

**Figure 3:**
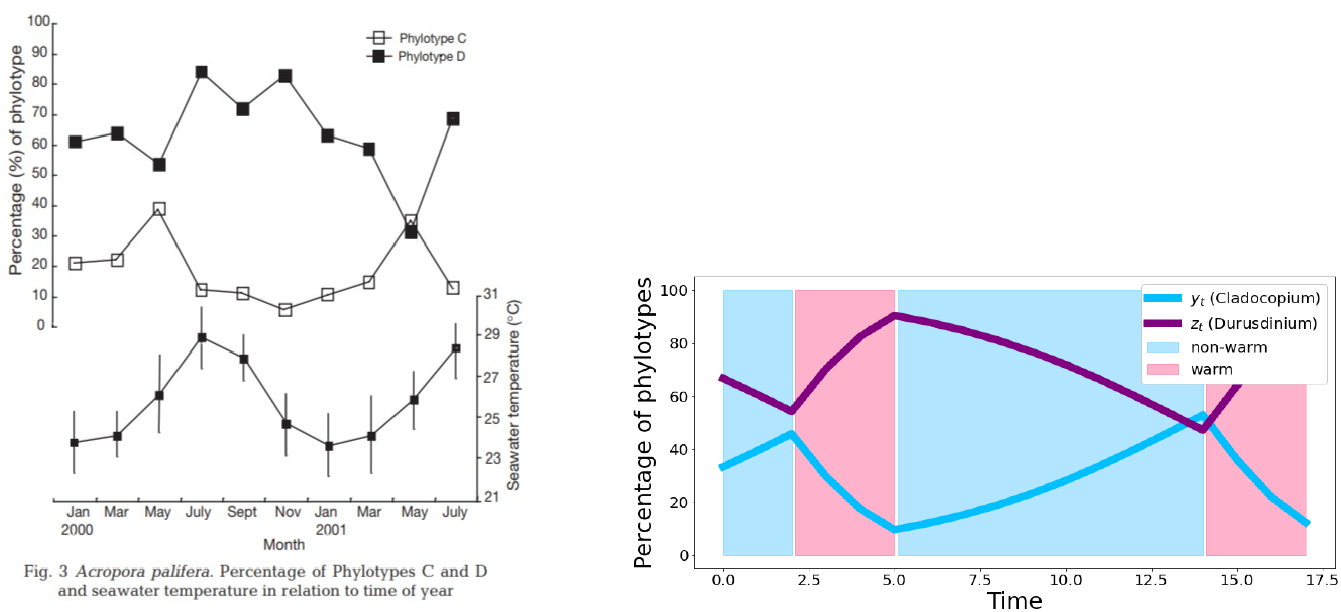
Comparison between observations from Chen et al. [2005] (Figure reproduced unchanged, CC) and the output of our model. The value of *θ* implements the different temperatures: cold for 1 (blue background) and warm for 0 (red background). The percentage of both symbionts changes according to the temperatures.

**Figure 4:**
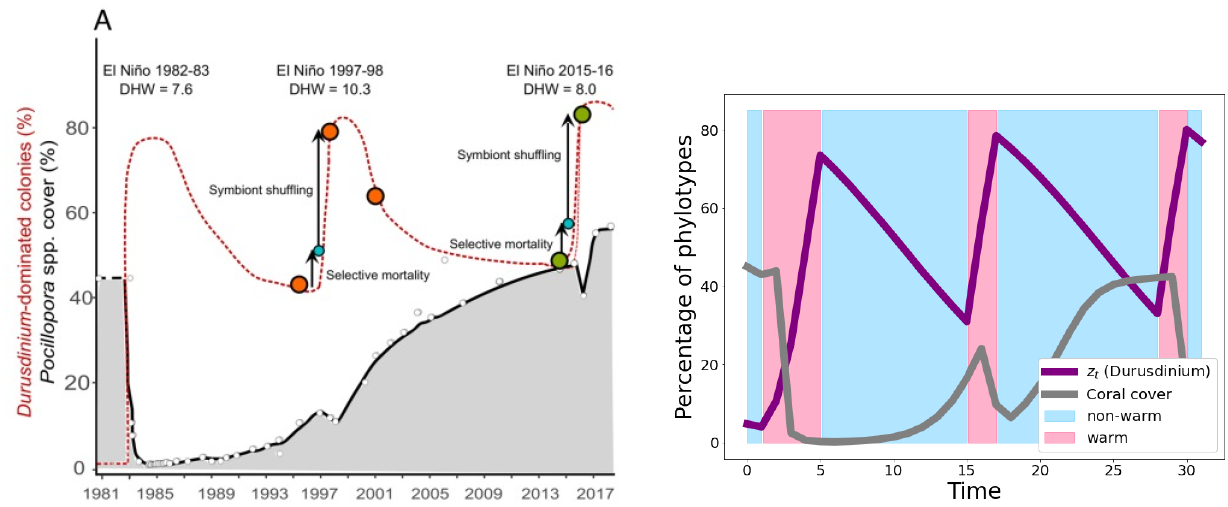
Comparison between observation from Palacio-Castro et al. [2023] (Figure reproduced unchanged with the kind permission of the authors, CC) and the output of our model. The model replicates the increase in the proportion of Durusdinium during warm periods.

In Figure 2, the brown color indicates El Niño, a warm period for corals. Our simulations (on the right) introduce a warm period for several time steps followed by a cold period. The simulation begins with a predominance of Cladocopium (*y*), the relatively more efficient algal species. The model accurately replicates the observed decrease in the total density of algae at the onset of the warm period and the shift in symbiont species from Cladocopium to Durusdinium during bleaching. During the subsequent cold period, the total density of algae remains relatively stable.

In Figure 3, we replicate the observations by Chen et al. [2005] (on the left). They measured the percentage of phylotype *C* and phylotype *D* over a year and a half (on the top), along with the corresponding seawater temperature (on the bottom). In our model (on the right), we roughly mimic the temperature trends using two temperature categories: cold (blue background) and warm (red background). The pattern used is cold, warm, cold, warm, to roughly match the observed temperature fluctuations by Chen et al. [2005]. The simulations start with approximately three times more phylotype *D* than phylotype *C*, mirroring the initial observations made by Chen et al. [2005]. The model predicts a slight shift from phylotype *D* to phylotype *C* during the first stage, followed by a quicker reversal during the second, warmer period. After the summer, the composition slowly shifts back to phylotype *C*, with another reversal during the last warm period of the observations.

In Figure 4, we compare the model’s output with observations from Palacio-Castro et al. [2023], depicting the proportion of the resilient algal species Durusdinium under different temperature regimes. The left panel shows the empirical data and hypothetical trajectory, while the right panel illustrates the model’s simulation results. During warm periods (red background), the proportion of Durusdinium increases, reflecting its temperature resilience. This shift in algal composition aligns well with the observed data. We also attempt to reproduce the coral cover measures using the following dynamics, with a scaling parameter *r*_*n*_ = 2.5. ^5^

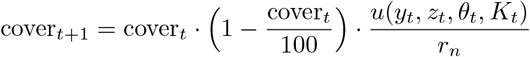

### Distribution of algae species

Most of the time, for any specific environment and coral species, one algal species dominates the others [de Souza et al., 2022, LaJeunesse et al., 2010]. The remaining algal species persist but at substantially lower densities[Silverstein et al., 2012, Mieog et al., 2007, Ziegler et al., 2018, Quigley et al., 2014, Bay et al., 2016, Buzzoni et al., 2023]. In our model, the composition of algae results from the competition between different algal species, influenced by their distinct growth rates across varying temperature conditions. The ratio of the two algal populations at time *T* is *η* to the power of the number of time steps where the environment was cold *θ*_*t*_ = 1, multiplied by *λ* to the power of the number of time steps where the environment was warm *θ*_*t*_ = 0, multiplied by the initial ratio of the two algal populations.

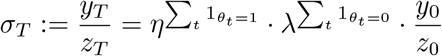

Specifically, the symbiont composition follows an exponential function of the geometric average of growth rates. This implies that if the environment remains stable for a sufficiently long period 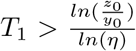 in the non-warm environment and 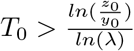 in the warm environment) one algal species will eventually out-compete the other [Little et al., 2004]. Hence, in environments with limited volatility, one algal species will typically dominate. Our model clarifies the impact of temperature on the diversity of algal species in coral symbiosis. According to our predictions, a prolonged period without environmental shifts allows one species to out-compete and dominate the others.

### Composition change of algae species

It has been observed that algal composition changes with increasing temperatures [Jones et al., 2008, Silverstein et al., 2015, Baker et al., 2004, Hsu et al., 2012]. Elevated temperatures can cause a shift in algal populations from the efficient Cladocopium to the temperature-resilient Durusdinium Buzzoni et al. [2023]. Interestingly, this is not always the case, as populations can revert to their original algal species[de Souza et al., 2022], a phenomenon known as *shuffle back* [Thornhill et al., 2006]. The drivers of a change from efficient algae to resilient algae are the duration of warm temperature [Cunning et al., 2015], and the repetition of warm events [Quigley et al., 2022]. Our model predicts *shuffle back* as a function of the environmental regime, including the duration of time spent in the same environment, and the initial composition of algae species. Once more, the symbiont composition follows an exponential trend with the geometric average of growth rates.

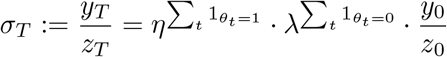

This exponential dynamic clarifies why short warm events may not change algal composition significantly (with the initial ratio being the dominant factor) [Bay et al., 2016], whereas prolonged warm events lead to a significant shuffle in algal composition (the exponential dynamic will be then decisive) [Cunning et al., 2018].

### Density population of algae species

Venn et al. [2008] showed that changes in the composition of algal species do not necessarily require bleaching. Consistent with this, our model demonstrates that the composition can shift without a significant decline in *K*_*t*_ (which is defined as bleaching). The surrounding temperature environment can gradually alter the algal composition without causing a major reduction in the total algal abundance. Furthermore, the model illustrates that such adjustments in symbiont composition are part of a normal adaptive process rather than an indication of stress. This aligns with the findings of Venn et al. [2008], reinforcing the idea that corals possess inherent flexibility in their symbiotic relationships, enabling them to adapt smoothly to environmental changes. This allows the coral to optimize its energy intake and maintain health without the dramatic loss of symbionts that characterizes bleaching events.

The experiments conducted by Cui et al. [2023] demonstrated that nutrient supplementation impacts symbiont density. Specifically, the provision of additional carbohydrates significantly reduces symbiont density, while ammonium promotes symbiont proliferation. On one hand, ammonium is a nitrogen source that is crucial for algal proliferation. When ammonium is provided, it enhances the growth rates of both Cladocopium and Durusdinium, promoting overall symbiont density. On the other hand, when additional carbohydrates are supplied, the coral does not anymore need to maintain a costly relationship with most of the algal population, since it is getting some carbohydrates anyway. Then, the coral will provide less nutrients to the population of algae, and the algae population will decrease. This nutrient-dependent regulation aligns with our model’s prediction that the coral optimizes nutrient allocation (*K*_*t*_) to balance the trade-offs between energy acquisition and algal population symbiosis cost.

The population densities of algae within coral symbioses are dynamic and are significantly influenced by environmental conditions. Seasonal changes, in particular, play a role in these fluctuations. Studies have documented these patterns, highlighting how varying temperatures throughout the year affect algal populations [Hoegh-Guldberg and Hinde, 1986, Stimson, 1997, Fagoonee et al., 1999, Fitt et al., 2000, Chen et al., 2005]. Our model captures these dynamics by considering how the coral regulates nutrient supply to different algal species in response to temperature changes. Specifically, the predicted optimal nutrient provision by the coral to the algal population changes with the temperature *θ*_*t*_. Observations by Buddemeier and Fautin [1993] indicate that in the absence of stress (i.e., in a consistently cool environment), the total population of algae remains notably stable. Within the framework of our model, this stability can be visually observed in the time series and can be verified formally. Firstly, Figure 5 shows the densities of Cladocopium (*C*) and Durusdinium (*D*), along with their sum (in black), representing the total algal population, which corresponds to *K*_*t*_, the amount of nutrients provided by corals. In a stable, cool environment (when *θ* = 1, depicted in blue), the model predicts an almost constant total algal population, even as the composition of algae changes. Secondly, we can analyze this stability formally. Recall that *K*_*t*_ is characterized as the total population of algae, with the optimal value given by

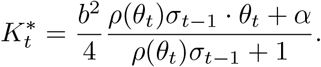

In a non-warm environment, this simplifies to

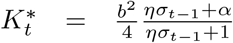

When *α* ≈ 1, the total population of algae 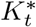 remains approximately constant and independent of the relative density of algal species *σ*:

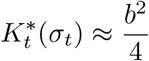

To make this statement more precise, consider the impact of the difference 1 − *α* on *K*_*t*_:

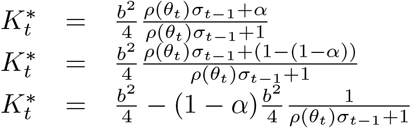

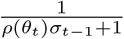 is smaller than 1 and we have that as 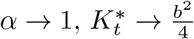

**Figure 5:**
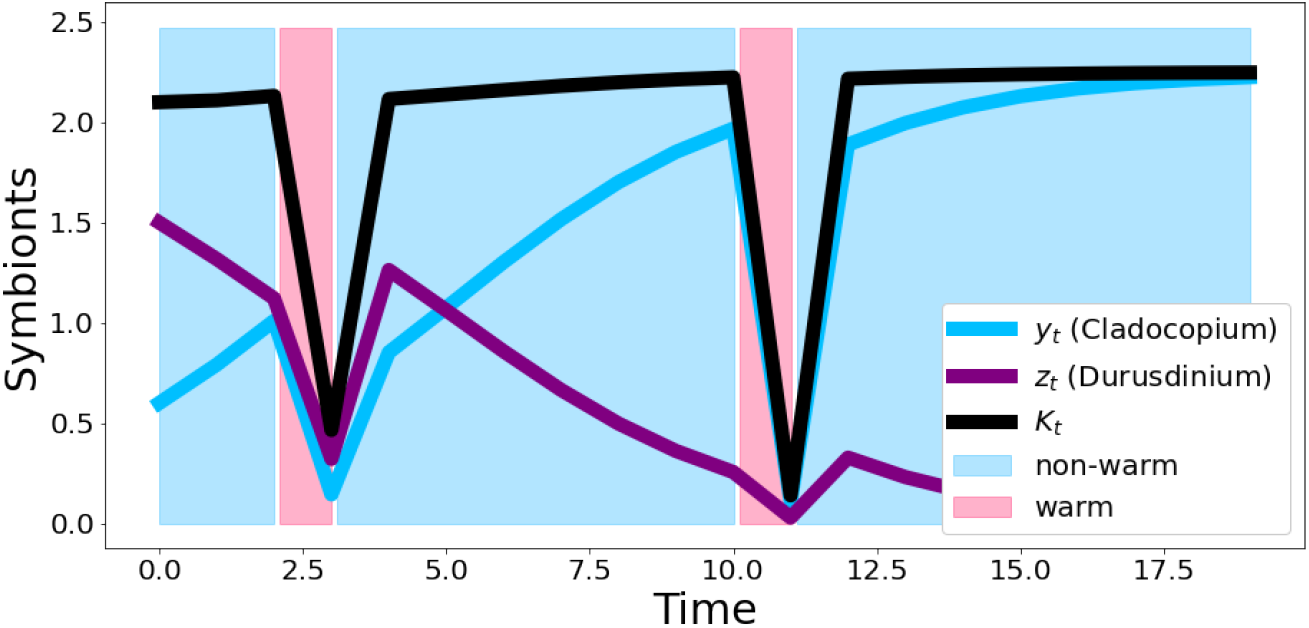
Model output showing that *K*_*t*_, the total population of algae, remains nearly constant in a stable environment.

**Figure 6:**
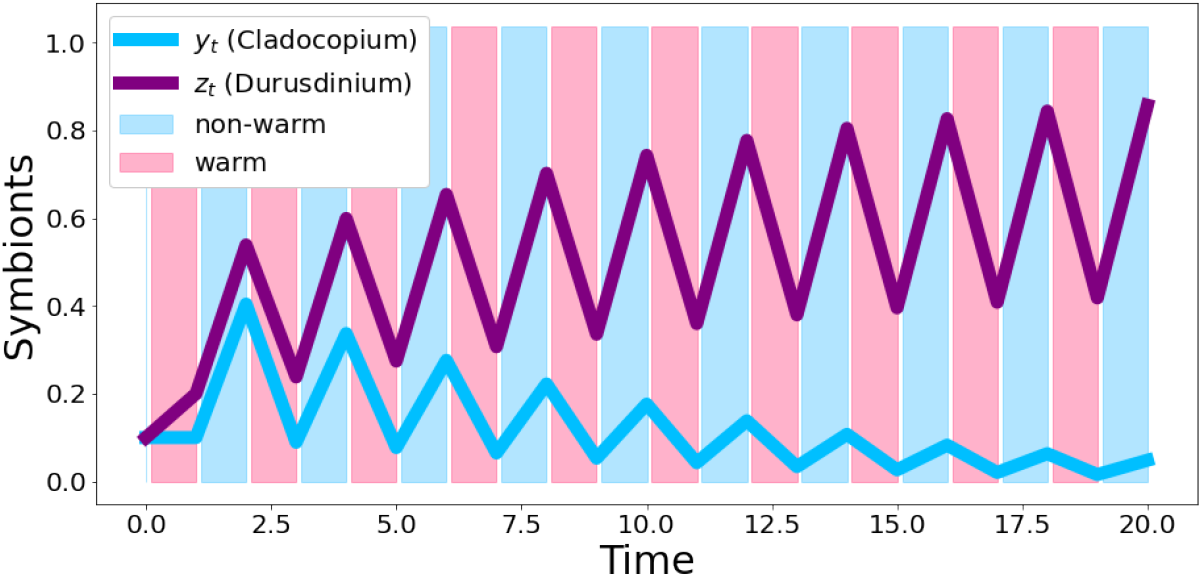
Plot of the model for *η* = 1.5 and *λ* = 0.5. The resilient algae outcompete the efficient ones over the long term.

### Bleaching severity

It has been observed that bleaching is less severe when the proportion of resilient algae is higher [Palacio-Castro et al., 2023, Fabricius et al., 2004, Douglas, 2003, Quigley et al., 2020, Baker, 2004]. In particular, Cunning et al. [2015] found that the severity of bleaching diminishes proportionally to the relative abundance of the resilient symbiont. Our model defines bleaching severity as the ratio of the total algae population in non-warm environments to the total population in warm environments. This ratio quantifies the reduction in the total population when the environment becomes warm. Formally, it is expressed as:

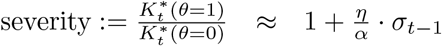

According to this definition, severity is an affine function of the previous period’s algae composition. Assuming that *λ* and *η* are not significantly different, the severity of bleaching is given by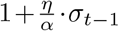, where *σ*_*t*−1_ is the ratio of the density of *y*_*t*−1_ to *z*_*t*−1_. Since 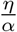 *α* is strictly positive, increasing the ratio *σ*_*t*−1_ will increase severity. This provides a rough estimate of how the proportion of each algal species influences the severity of bleaching. Our model aligns with empirical observations, indicating that a higher proportion of temperature-resilient algae can mitigate bleaching severity. This insight underscores the critical role of symbiont composition in enhancing the resilience of corals to thermal stress.

## 5 Generalization for any temperatures

In this section, we extend the model to accommodate an arbitrary number of temperature environments. The algae dynamics are as follows:

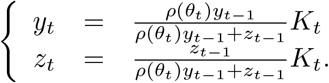

The coral’s instantaneous payoff remains *u*(*y*_*t*_, *z*_*t*_, *θ*_*t*_, *K*_*t*_) = *b*(*y*_*t*_, *z*_*t*_, *θ*_*t*_)−*c*(*θ*_*t*_, *K*_*t*_), but with a slightly different benefit function 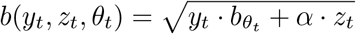 and the feeding cost still given by *c*(*θ*_*t*_, *K*_*t*_) = *K*_*t*_. The optimal quantity of nutrients the coral should provide to algae, for each environment, is now expressed as:

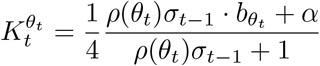

In this generalized model, both the proportion of each algal and the total algae densities change continuously with the environment Fautin and Buddemeier [2004]. This aligns with observations that the composition of symbionts reacts to small changes in the environment [de Souza et al., 2022].

Resilient algae are often found in more variable environments [Baker et al., 2013a, Carballo-Bolaños et al., 2019, Oliver and Palumbi, 2011]. It has been shown that high-frequency temperature variability significantly reduces bleaching [Safaie et al., 2018, Sully et al., 2019, Carilli et al., 2012]. Conversely, bleaching is more prevalent where thermal-stress anomalies are more frequent [Sully et al., 2019]. Additionally, a bimodal temperature distribution has been shown to decrease bleaching prevalence [McClanahan et al., 2019]. According to our model, temperature variability affects the composition of algae differently compared to a stable temperature environment. More precisely, resilient symbionts can slowly out-compete efficient ones if temperatures fluctuate sufficiently, which would not be possible under constant temperature conditions. Mathematically, since the geometric average is always lower than the arithmetic average, temperature variability (within a low enough range) provides a relative advantage to resilient algae species, thereby increasing coral resilience.

However, if temperatures increase excessively (such as in the IPCC’s *business as usual* scenarios [Pörtner et al., 2019]), even resilient symbionts may not survive, leading to dramatic declines in coral populations. We hypothesize a non-linear relationship between temperature increases and coral survival. While a small temperature rise might only slightly reduce coral survival [Rowan, 2004] due to trade-offs between temperature-sensitive and temperature-tolerant algae, higher temperatures could be significantly more detrimental. Logan et al. [2014] estimate that symbiont shuffling alone will delay most bleaching events by approximately 10 years under low-emission scenarios, whereas Palacio-Castro et al. [2023] estimate a delay of 25-35 years in the eastern tropical Pacific.

## 6 General discussion

Coral-algae symbiosis operates near its thermal threshold, tolerating temperatures close to their optimal range. This threshold is influenced by factors such as depth [Iglesias-Prieto et al., 2004], location, and local conditions [Berkelmans, 2002]. While living close to this thermal limit might seem precarious, corals have developed a strategic advantage through their ability to reshuffle symbionts. This adaptability allows corals to optimize energy intake and maintain health even under stress, highlighting the potential adaptive benefits of bleaching in certain contexts [Douglas, 2003]. The capacity for symbiont reshuffling is crucial for the long-term survival of corals, helping to reconcile the paradox that coral reefs appear sensitive to short-term environmental perturbations but remain robust over geological timescales [Fautin and Buddemeier, 2004, Buddemeier et al., 2004]. However, this resilience has its limits, especially as climate change drives environmental conditions beyond historical norms [Schoepf et al., 2023]. Catastrophic bleaching may represent the end of a spectrum that includes natural fluctuations in symbiont populations over seasonal timescales. Such bleaching events, while dramatic, are not always necessary for adaptive or acclimatory responses in coral ecosystems [Buddemeier et al., 2004]. In environments with slowly and weakly changing temperatures, corals can effectively acclimate, maintaining their symbiotic relationships and overall resilience.

The temperature pattern is crucial for predicting future coral reef survival [Cunning, 2021]. Our model shows that a long period without warm temper-atures 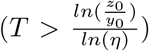 can lead to a shift back to the less resilient algal species, making corals more vulnerable to future warm periods. Thus, infrequent temperature warming could be more dangerous than frequent warming, as corals better adapt to consistent thermal stress. This characteristic has also been explained in the model of Ware et al. [1996]. Our model suggests that effective symbiont shuffling enhances coral resilience within a specific range of temperature increases. However, once temperatures exceed the threshold of resilience for the algae, shuffling no longer provides additional benefits and can result in coral declines. For more detailed projections on coral bleaching, refer to Logan et al. [2014] and Palacio-Castro et al. [2023].

While this study focuses on coral-algae symbiosis, the principles could extend to coral-microbiome interactions [Ziegler et al., 2017, Voolstra and Ziegler, 2020, Maire et al., 2023a,b, Bourne et al., 2016, Torda et al., 2017, Doering et al., 2021, van Oppen and Blackall, 2019, Marangon et al., 2021, Tran, 2022, Wei et al., 2024, Gardner et al., 2023, Voolstra et al., 2024]. Coral microbiomes, like algal symbionts, are diverse and can buffer against environmental fluctuations. Nutrients or signals provided by corals could regulate these microbial populations, contributing to overall coral resilience [Rädecker et al., 2015]. For a model incorporating coral microbiomes, see Lima et al. [2020]. Likewise, studies such as Camp et al. [2020] and Chen et al. [2024] highlight interactions between algae and fungi under thermal stress. Similarly, the sole environmental variable accounted for in the model is sea temperature; nevertheless, a comparable phenomenon may occur with light.

Acclimation to higher temperatures via symbiont shuffling might reduce resistance to other disturbances. Given the trade-off between efficiency and resilience, there may also be trade-offs between resilience to different disturbances, such as temperature and acidity [Anthony et al., 2011, Van der Zande et al., 2020], or even combined effects of temperature, acidity, and diseases [Ateweberhan et al., 2013]. Climate change might not only reduce the efficiency of the algal symbiosis but also lower resilience to other stressors. Reviews on coral reefs under climate change and ocean acidification can be found in Anthony [2016] and on multiple stressor interactions in Ban et al. [2014]. Considering multiple stressors is crucial, as emphasized by Pendleton et al. [2016] and further discussed in Bieg et al. [2024]. Local stressors have shown varying effects on corals [Pancrazi et al., 2020, Cannon et al., 2023, Prazeres et al., 2017, Jones and Gilliam, 2024, Baum et al., 2023], with some studies finding no significant impact [Johnson et al., 2022]. For a comprehensive review, see Dutra et al. [2021]. Models that include local stressors can be found in Hafezi et al. [2020] and Gurney et al. [2013]. Integrating considerations of ocean acidification and warming with symbiont dynamics will be essential for future research [Hoadley et al., 2015, Grottoli et al., 2018].

## 7 Conclusion

In this paper, we presented a model that captures the intricate dynamics of coral-algae symbiosis in response to fluctuating environmental conditions. We developed a model wherein a coral optimizes the quantity of nutrients provided to its algal symbionts, specifically two distinct species: one that is relatively more efficient at carbon fixation and another that is more resilient to temperature increases. These species compete for the nutrients supplied by the coral. Our model explains the phenomenon of bleaching as an adaptive process within coral symbiosis. This stylized model enables us to analytically express several key quantities. We find that the severity of bleaching can be predicted as a function of the current temperature and the relative proportions of each algal species, independent of the absolute density of the algae populations. Furthermore, the future composition of algal species is anticipated to be proportional to the current composition, adjusted by the geometric average of the growth rates associated with the different environments. This allows us to hypothesize about the limits of coral adaptation through symbiont change when faced with temperatures that exceed the tolerance of even the resilient algal species. With only a few parameters, our analytical model provides explanations for various observed phenomena, including the responses of total algal population density under both constant and variable environmental conditions. It also accounts for specific features noted under different temperature regimes, such as the *shuffle back* phenomenon during brief warming periods and the dominance of resilient algal species in environments with high-temperature variability.

The analytical expressions derived from our model facilitate potential statistical testing, enabling the validation of the model’s predictions against empirical data. Additionally, this model can be applied in conjunction with climate change projections to assess the potential survival of coral reefs in the coming decades. By integrating other factors, such as ocean acidity and biodiversity loss, this model could serve as a foundational framework for a more comprehensive understanding of coral ecosystem dynamics in the face of global environmental change.

### A Typical time scale to shuffle back to the efficient genera

We want to prove that the typical time that it takes to achieve a majority of the efficient genera (*y*) when the environment is always in the non-warm state is 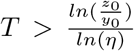. We here define as the typical time that it takes to achieve a majority of the efficient genera (*y*) as the first time when *y*_*t*_ *> z*_*t*_, which is equivalent to *σ*_*t*_ *>* 1 (by definition). Recall that we have

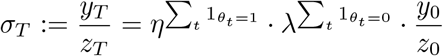

Since we consider a succession of only non-warm states, we have

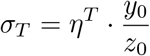

Then,

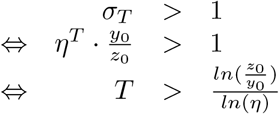

Similarly, we can prove that the typical time that it takes to shuffle to the resilient genera when the environment is always in a warm environment is 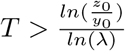.

### B ptimum amount of nitrogen for any temperatures

We here generalize previous results for any temperature. The instantaneous coral payoff is still *u*(*y*_*t*_, *z*_*t*_, *θ*_*t*_, *K*_*t*_) = *b*(*y*_*t*_, *z*_*t*_, *θ*_*t*_) − *c*(*θ*_*t*_,*K*_*t*_), with benefit function different as before 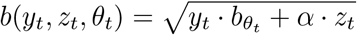, and the feeding cost is still the same *c*(*θ*_*t*_, *K*_*t*_) = *K*_*t*_.

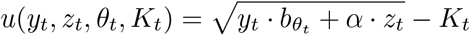

The dynamics of algae are still,

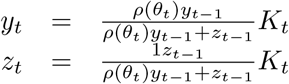

We can rewrite, again, the expression of the payoff, as a function of *θ*_*t*_ and *K*_*t*_. Again, recall that by definition, 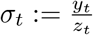.

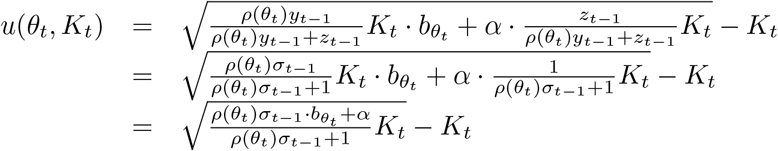

Consider the derivative with respect to the control parameter *K*_*t*_.

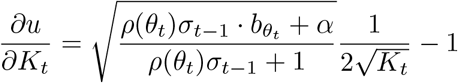

The derivative is null only at one point,

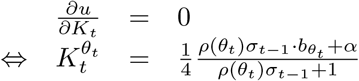

The optimum quantity of nutrients the coral should provide to the population of algae is then:

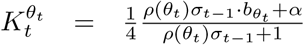

### C Payoff generalization

In this section, we want to show that there is exactly one optimum for a broader definition of benefit and cost function. We use the following definition of benefit and cost.

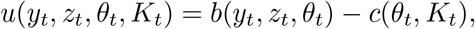

With

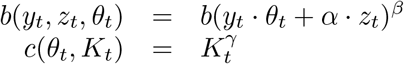

The dynamic of algae is still the same,

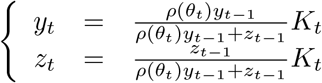

We can rewrite, again, the expression of the payoff, as a function of *θ*_*t*_ and *K*_*t*_.

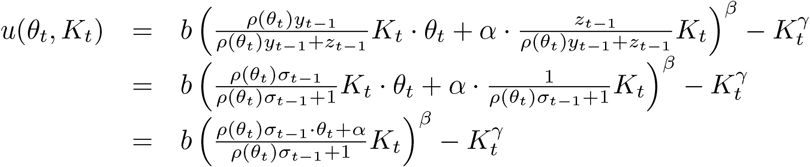

The first derivative,

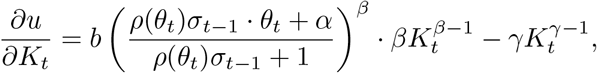

is null only at one point in the interior, i.e., in (0, ∞), for *β < γ*.

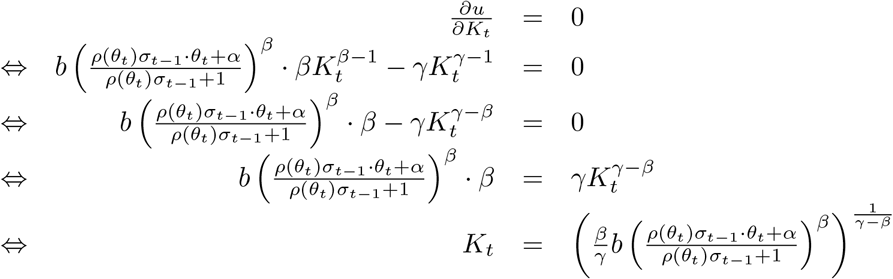

### D Shuffling and switching

In this appendix, we want to show that, in general, the dynamic of alga is driven by the growth of alga already present in the symbiosis. While it has been shown that shuffling (a change in the proportion of species of algae already present with the host) is more significant than switching (the establishment of symbiosis with new species of algae) [Berkelmans and Van Oppen, 2006, Jones et al., 2008], corals have still the possibility to establish new symbiosis [Boulotte et al., 2016]. Moreover, according to Sampayo et al. [2008] the main driver of composition change is the differential mortality of algae. We propose a variation of the model described earlier to consider the possibility for the coral to establish new symbiosis. We note *τ* the quantity of new symbiont.

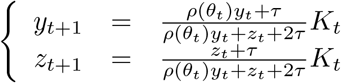

We can show that:

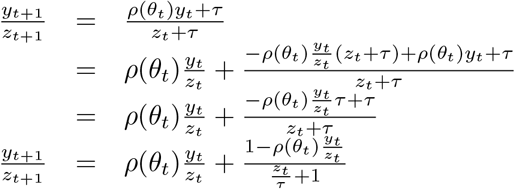

For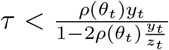, we have that shuffling as a larger influence than switching. In particular, if *τ* the quantity of new symbiosis is small enough, the dynamic of algae will not be significantly affected. This means that if we relax the assumption to allow both *shuffling* and *switching*, as long as the influx of new symbionts is sufficiently small (a quantity we can characterize), the dynamics of the algae are not dramatically impacted, and the previous results remain valid.

### E Non-myopic corals

In this section, we assume corals optimize their fitness in the long term and not only for the present time. We consider again two states of the environment *θ* ∈ {0, 1} with 0 the *bad* state and 1 the *good* state. However, now the two states occur according to a Markov model, with transition probabilities *p*_00_, *p*_01_, *p*_10_, and *p*_11_, where *p*_*ij*_ is the probality of a switch from state *i* ∈ {0, 1} to state *j* ∈ {0, 1}. The dynamics of the algae are given by

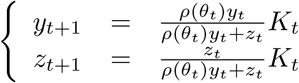

The overall payoff function is now

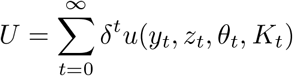

Both the benefit and cost functions remain unchanged. The optimal nutrient quantity for the two states can be determined as follows, though the solution has become somewhat more complex.

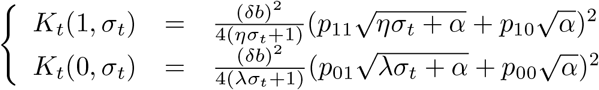

Notice that for *σ*_*t*_ → 0 the influence of the stochastic process is negligible on the optimum *K*_*t*_(1, .) and *K*_*t*_(0, .). This implies that if the population consists primarily of resilient algae, the optimal quantity of algae will remain consistent across all expected future states, as the resilient algae will perform equally well under varying conditions. Note also that the generalized solution here embeds the previous solution. Indeed, for *δ* → 1, *p*_11_ → 1 and *p*_00_ → 1 we have the same optimum (for the next step). Consider the relative change between the two different model results (*m* refers as the myopic –basic– model, and *nm* as the non-myopic model, presented here):

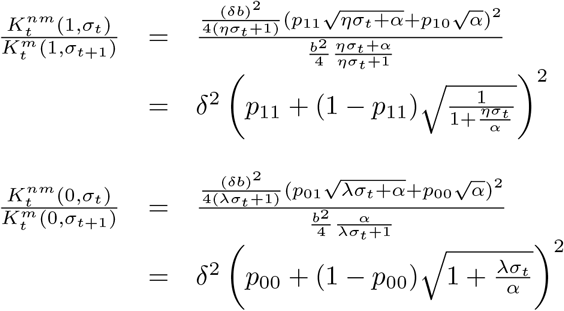

With this model, we find that the optimal 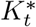 depends on the stochastic regime (i.e., *p*_*ij*_). If corals are indeed adapted to a specific stochastic regime (i.e., a specific value of *p*), moving corals from one sea to another with a different regime could result in maladaptation to the new stochastic environment. Therefore, translocating corals between seas requires careful consideration of various environmental factors, especially temperature conditions, to ensure their well-being.

## Footnotes

1 Discrete time makes it relatively simple to model a coral who makes decisions now (at time *t*) to affect the future (at time *t* + 1).

2 This generality is possible due to our assumption that the coral is myopic, focusing on immediate future states without long-term foresight. Refer to Appendix E for the scenario involving a forward-looking coral.

3 Numerous algal species have been identified within corals [LaJeunesse et al., 2018, Yorifuji et al., 2021]. The restriction to two key types, one efficient and the other more resilient, is made for simplicity and tractability. We believe that this restriction is sufficient to capture the main mechanism behind coral bleaching.

4 Additionally, Fagoonee et al. [1999] demonstrated the ability of corals to regulate algal populations. Models by Raharinirina et al. [2017, 2022] also incorporate the concept that *corals are in control of the symbiotic relationship*.

5 This basic dynamic is introduced to illustrate the relevance of the payoff definition, rather than to explicitly model coral cover.

